# Nanobodies as therapies for loss-of-function misfolding diseases

**DOI:** 10.1101/2025.04.12.648492

**Authors:** Atanasio Gómez-Mulas, Mirco Dindo, Juan Luis Pacheco-García, Silvia Grotelli, Mario Cano-Muñoz, Pavla Vankova, Dmitry S. Loginov, Eduardo Salido, Petr Man, Francisco Conejero-Lara, Barbara Cellini, Angel L. Pey

**Affiliations:** Departamento de Química Física, Facultad de Ciencias, Universidad de Granada, 18071 Granada, Spain; Department of Medicine and Surgery, University of Perugia; Institute of Biotechnology - BioCeV, Academy of Sciences of the Czech Republic, Vestec, Czech Republic; Institute of Microbiology - BioCeV, Academy of Sciences of the Czech Republic, Vestec, Czech Republic; Center for Rare Diseases (CIBERER), Hospital Universitario de Canarias, Universidad de la Laguna, 38320 Tenerife, Spain; Departamento de Química Física, Instituto de Biotecnología y Unidad de Excelencia de Química Aplicada a Biomedicina y Medioambiente (UEQ), Facultad de Ciencias, Universidad de Granada, 18071 Granada, Spain

**Keywords:** Hyperoxaluria, Protein misfolding, Inborn error of metabolism, Therapeutics, Nanobodies

## Abstract

Misfolding diseases that result in loss of function represent a considerable burden for both individuals and society. Primary hyperoxaluria type 1 (PH1) is a rare genetic disorder caused by mutations in the alanine:glyoxylate aminotransferase 1 (AGT) enzyme. The underlying molecular mechanisms causing PH1 are associated with protein misfolding (enhanced aggregation and mitochondrial mistargeting). The main therapeutic approach to increase patients’ lifespan and quality of life is a double kidney and liver transplantation. Alternative treatments such as gene and enzyme replacement and pharmacological chaperones are currently being introduced, but other alternatives are necessary. In this work, we developed and characterized a novel biotechnological approach using six single-domain nanobodies (NB-AGT-1 to -6) as potential therapeutics for PH1 misfolding. We show that NB-AGTs are very stable proteins and bind to pathogenic and non-pathogenic variants of AGT with extreme affinities (with K_d_ values from low nM to low pM). Structural studies showed that NB-AGTs bind to different epitopes of AGT with selectivity for different AGT variants. Experiments in cellular PH1 models showed that internalization of engineered NB-AGT-3 enhanced the specific activity of disease-associated variants. Overall, we show that NBs are a novel and promising approach to treat PH1 and other loss-of-function misfolding diseases.

## 1. Introduction

Protein folding in vivo is a complex process by which a polypeptide typically searches for the native state populating multiple conformations and interacting with the protein homeostasis network [1,2]. This network operates by helping proteins to fold into their native state, preventing aggregation or targeting for degradation [1,3]. Those protein conformations populated during folding or existing in the dynamic and heterogeneous native state ensemble are associated with protein aggregation and enhanced degradation are typically called *misfolded* states [1,2,4,5]. Mutations in the genome of somatic and germinal cell lines often cause single amino acid changes (missense variations) that enhance protein misfolding, and may lead to human disease [5–9]. Misfolding diseases are typically classified into two large groups. The loss-of-function diseases in which a missense variation in the sequence decreases the intracellular protein function (usually promoting protein degradation or the formation of inactive aggregates). Whereas in gain-of-function diseases the variation enhances a toxic function (for instance, the formation of toxic aggregates) [2,5,7,9–11]. In the context of protein folding and misfolding in human diseases, a simple approach to overcome misfolding is to target the stability of native proteins using (often small) natural or unnatural ligands that preferentially bind to and stabilize it [5,7,8,12–19]. Some limitations of these approaches concerning small ligand bioavailability and limited operational binding affinity (and therefore stabilization of the native state) might be overcame using high-affinity selective binders such as specific nanobodies [20–26].

Primary hyperoxaluria type I (PH1) is a rare inborn metabolic disease caused by a deficit in the peroxisomal alanine:glyoxylate aminotransferase enzyme 1 (AGT), causing oxalate overproduction and renal failure [27]. AGT exists in two versions in human population, a most common (major haplotype) wild-type (AGT-WT) form and a polymorphic (minor) variant (AGT-LM, containing two amino acid substitutions, P11L and I340M). The latter is not pathogenic by itself, but it predisposes to develop the disease in synergy with additional rare mutations [8,27]. Interestingly, over 150 mutations (half of them on the minor allele) have been reported, but two of them (LM-G170R and LM-I244T) account for about 40% of the alleles in patients [8,27,28]. Interestingly, these two mutations have been classified as operating through different mechanisms, LM-G170R mistargeting the enzyme to the mitochondria where it is metabolically useless, and LM-I244T that enhances protein aggregation and inactivation in peroxisomes [27–31]. However, PH1-associated misfolding leading to aggregation or mistargeting might be intertwined mechanisms in structural and energetic terms [32,33].

Currently, the most widely used therapeutic approach to extend the life-span and to increase the quality of life of PH1 patients is a double liver-kidney transplantation, which is obviously difficult and problematic [27]. Other therapeutic approaches are under investigation, some of them targeting the instability and misfolding of disease-causing mutants, such as supplementation with precursors of AGT cofactor (pyridoxal 5′-phosphate, PLP) and the use of pharmacological chaperones [14,31,34]. Consequently, we need to find alternative ways to stabilize mutant variants of AGT, particularly with tight binding to AGT variants and pharmaceutical stability, which applying the mass action law should stabilize the native state and may correct misfolding.

In this work, we present an alternative treatment for PH1 using nanobodies (NB) as native state stabilizers of AGT (referred to as NB-AGTs through the manuscript). NB are single-domain proteins derived from the heavy-chain antibodies of camelids, with a great potential as therapeutic agents [20–26]. NBs stood out as pharmacological agents due to their very high stability and tight binding to their target proteins [20,24,26,35,36]. We have developed 6 NB-AGTs upon llama immunization, characterized their interaction with different pathogenic and non-pathogenic variants of AGT, and provided evidence that their therapeutic potential is high using cell models of PH1 by adding to the NB-AGTs sequence a strong cell-penetrating peptide (CPP)[37].

## 2. Materials and Methods

### 2.1. Generation of NB-AGTs

Nanobody generation was carried by carried out by NabGen Technologies (Marseille, France).

#### 2.1.1. Llama immunization

A llama was immunized with the purified AGT-LM protein dimer (purified as described in [31]). 500 µg of the AGT-LM protein were subcutaneously injected 3 times at 21 day intervals using incomplete and complete Freund’s adjuvant (ThermoFisher, Illkirch-Graffenstaden, France). A blood sample was collected aseptically 12 days after the last injection. Pre-immune and immune sera were collected at day 0 and day 54 (after immunization).

AGT-LM was adsorbed on a maxisorp plate (ThermoFisher, Illkirch-Graffenstaden, France). The immune response was analysed by ELISA by serum dilution 1/250 or 1/500 in PBS, using goat anti-llama-HRP conjugate (Bethyl A160-100P, Waltham, MA, USA). Absorbance was recorded in 96-well plates at 450 nm on a Safire 2 microplate reader (Tecan, Lyon, France).

#### 2.1.2. Library construction

The blood collected on day 54 was treated within less than 24 hours to avoid coagulation. PBMCs were separated by histopaque density gradient. mRNA was then extracted using the RNeasy Mini Kit from Qiagen (Les Ulis, France). cDNA was synthesized using the SuperScript IV First strand Synthesis System Invitrogen (ThermoFisher, Illkirch-Graffenstaden, France). The nanobody encoding reading frames from this cDNA were amplified by PCR and then cloned into a phage display vector with a Human influenza hemagglutinin (HA) tag.

#### 2.1.3. Panning by phage display

Enrichment for target specific antibodies against AGT was performed using the phage display technique with M13KO7 Helper phage (New England Biolabs, Courcouronnes, France). To immobilize the targets for selection by phage display, a specific adsorption onto the solid surface of a maxisorp plate was used.

#### 2.1.4. Selection by ELISA

Following two rounds of panning, 12 clones from round 1 and 12 clones from round 2 were picked and expressed in *E. coli*. Those clones were tested against AGT-LM and AGT-WT with an ELISA using a monoclonal anti-HA antibody produced in mice (Sigma-Aldrich, Lezennes, France) as a primary antibody and goat anti-mouse IgG/IgM HRP conjugated antibody (Millipore, Molsheim, France). Absorbances of wells were recorded at 450 nm on a Safire 2 microplate reader (Tecan, Lyon, France). Positive clones were sent for sequencing (Figure 1A).

**Figure 1.**
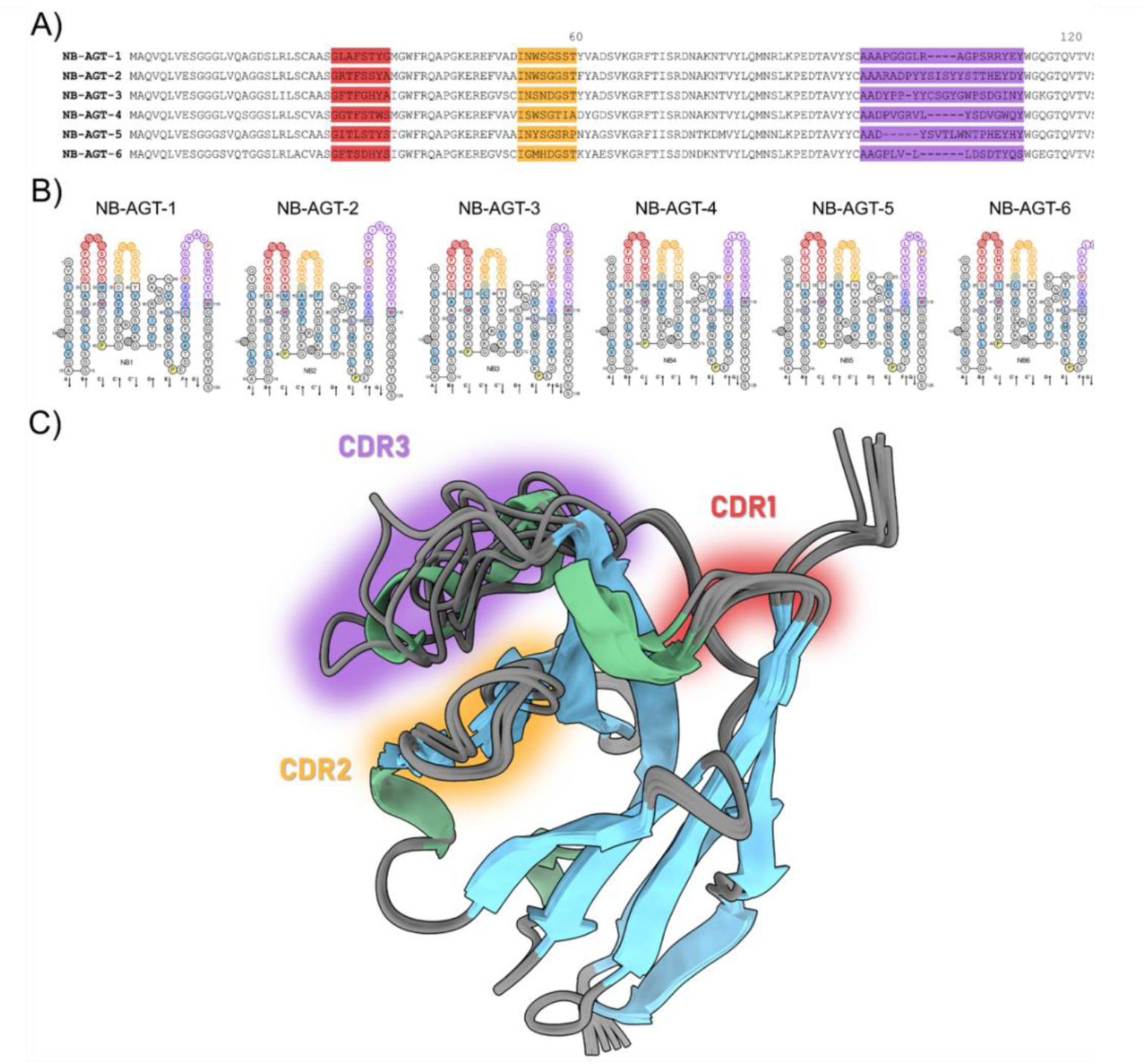
Sequence and structure of NB-AGT: **A)** Sequence alignment of NB-AGT variants generated using Clustal Omega ([70]; https://www.ebi.ac.uk/Tools/msa/clustalo/). Numbering corresponds to NB-AGT-1. Complementarity-determining regions (CDRs 1–3) are highlighted and are consistent with sequence alignment and structural models (loops in panel C). **B)** IMGT Collier de Perles on one layer. Amino acids are shown in the one-letter abbreviation. All proline (P) are shown online in yellow. IMGT anchors are in square. Positions with bold (online red) letters indicate the four conserved positions that are common to a V domain and to a C domain. The hydrophobic amino acids (hydropathy index with positive value: I, V, L, F, C, M, A) and tryptophan (W) found at a given position are shown with a blue background color. Arrows indicate the direction of the beta strands and their designations. **C)** Structural alignment of six NB-AGT variants. Structures were predicted using ColabFold v1.5.2–AlphaFold2 [44], https://colab.research.google.com/github/sokrypton/ColabFold/blob/main/AlphaFold2.ipynb#scrollTo=G4yBrceuFbf3) with default parameters and template_mode PDB100. Structural alignment was performed in PyMOL v2.5.2. Hypervariable loops in CDRs 1–3 were indicated. Pairwise RMSD was 0.32 ± 0.07 Å (NB-AGT-2 to 5 vs. NB-AGT-1). All six nanobodies are shown in ribbon and colored by secondary structure (α-helices colored in green and β-strands colored in light blue). CDR1 is colored in red, CDR2 in yellow and CDR3 in purple in all panels.

### 2.2. Protein expression and purification

*E. coli* BL21 (DE3) cells were transformed with the pET-24(+) vector containing the cDNA of each nanobody and carrying a C-terminal 6-His-tag (GenScript, Leiden, The Netherlands). 40 mL of LB preculture containing 30 µg·mL^-1^ kanamycin (LBK) was inoculated with transformed cells and grown for 16 h at 37 °C. These cultures were diluted into 800 mL of LBK, grown at 37 °C for 3 h and nanobody expression was induced by the addition of 0.5 mM IPTG (isopropyl β-D-1-thiogalactopyranoside) and lasted for 6 h at 25 °C. Cells were harvested by centrifugation and frozen at -80 °C for 16 h. Cells were resuspended in binding buffer, BB (20 mM Na-phosphate, 300 mM NaCl, 50 mM imidazole, pH 7.4) plus 1 mM PMSF (phenylmethylsulfonyl fluoride) and sonicated in an ice bath. These extracts were centrifuged (20000 g, 30 min, 4 °C) and the supernatants were loaded into IMAC (immobilized-metal affinity chromatography, Cytiva, Marlborough, MA, USA) columns, washed with 40 bed volumes of BB and eluted in BB containing a final imidazole concentration of 500 mM. These eluates were immediately buffer exchanged using PD-10 columns to 50 mM HEPES-KOH (HEPES, 2-[4-(2-hydroxyethyl)piperazin-1-yl]ethanesulfonic acid) pH 7.4. Protein purity was analyzed by SDS-PAGE and those with some impurities were loaded onto a Superdex 75 16/60 size exclusion chromatography column (Cytiva, Barcelona, Spain) using 20 mM HEPES-NaOH, 200 mM NaCl pH 7.4 as mobile phase at 1 mL·min^-1^ flow rate. Proteins were stored at -80 °C after flash-freezing in liquid nitrogen. Purity was analyzed again by SDS-PAGE. Protein concentrations were measured spectrophotometrically using the following molar extinction coefficients (ε_280_, in M^-1^·cm^-1^) according to their primary sequence: NB-AGT-1.- 25565; NB-AGT-2.- 33015; NB-AGT- 3.- 33140; NB-AGT-4.- 36565; NB-AGT-5.- 30035; NB-AGT-6.- 20065. The purity and molecular weight of each nanobody were further checked by high performance liquid chromatography coupled to electrospray ionization mass spectrometry (HPLC/ESI-MS) by the High-resolution mass spectrometry unit, Centro de Instrumentación Científica (University of Granada) in a WATERS LCT Premier XE instrument equipped with a time-of-flight (TOF) analyzer. The purity and integrity of purified NB-AGTs can be seen in Figure S1. The biophysical characterization of NB-AGT conformation is described in the Supplementary information. NB-AGT for experiments with cell cultures, we used the same construct that the one used for NB-AGT (CPP-NB-AGT) but adding a potent cell penetrating peptide (KRKKKGKGLGKKRDPCLRKYK) followed by a linker region (GSGSGS) to avoid interactions between the NB-AGT and the CPP (GenScript, Leiden, The Netherlands). Expression and purification of CPP-NB-AGTs were performed following the protocol described above for NB-AGTs.

### 2.3. Differential scanning calorimetry (DSC)

DSC experiments were carried out in MicroCal PEAQ-DSC microcalorimeter equipped with an autosampler (Malvern Panalytical, Malvern, UK). Scans were run from 5 to 130 °C at a scan rate of 120 °C·h^-1^. The experiments were done in 20 mM HEPES-NaOH, 200 mM NaCl pH 7.4. Protein concentration was 5-40 µM for the nanobodies and 5 µM (in monomer) for the AGT LM in the presence of 25 µM of pyridoxal 5′-phosphate (PLP, Merck, Madrid, Spain). Instrumental baselines were recorded before each experiment with both cells filled with buffer and subtracted from the experimental thermograms of the protein samples. To evaluate the effect of NB-AGT binding on the stability of AGT LM we subtracted the signal arising from a NB-AGT at a given concentration from the signal observed in AGT LM with the same concentration of NB-AGT, and by fitting this difference to a two-state irreversible model as described previously for AGT LM [30]. To determine the stability of NB-AGT, 20 µM NB-AGT solutions were used and DSC scans were carried out as described above for the complexes and fitted using a pseudo two-state reversible unfolding model

### 2.4. Surface Plasmon Resonance (SPR)

Binding affinity for the interaction between NB-AGTs and AGT variants was evaluated using a Biacore T200 Surface Plasmon Resonance instrument (Cytiva, Pensacola, FL, USA) essentially as described [38,39]. All six NB-AGTs were individually and covalently immobilized on NTA chips aiming for reaching 500 response units (RUs). For this, we first determined the amount of time required for each variant to reach this level of response. To capture the His-tagged NB-AGTs, the nitrilotriacetic acid (NTA) chip was loaded using a NiCl_2_ 0.5 M solution and 100 - 200 nM nanobody samples in a HBS-P+ buffer (10 mM HEPES, 150 mM NaCl, 0.05% v/v Tween20, Cytiva, Pensacola, FL, USA) passed under a flow of 5 µL/min. Once these times were determined, the immobilization procedure required activation of the NTA matrix with the same 0.5 M NiCl_2_ solution and activation of the carboxyl groups of the same matrix with a 1:1 mixture of N-ethyl-N-(3-diethylamino-propyl)-carbodiimide (EDC) and N-hydroxysuccinimide (NHS). Nanobodies followed for the time previously determined under a 5 µL/min flow and ethanolamine blocked the remaining activated carboxyl groups, ending the immobilization step. Each NTA chip has 4 flow cells in which, the first was activated with the EDC/NHS mixture and then blocked with ethanolamine without ligand ever flowing through so as to function as a reference for the others. Experiments were carried out at 25°C.

Affinity constants (*K*_a_; their inverse are the dissociation constants, *K*_d_, see Table 1) as well as the kinetic association (*k*_on_) and dissociation constants (*k*_off_) were determined by performing Sigle Cycle Kinetics (SCK) assays. In this case, serial dilutions of the different AGT variants in HBS-EP+ (10 mM HEPES, 150 mM NaCl, 3 mM EDTA, 0.05% v/v Tween20), with concentrations ranging 0.74-60 nM, were injected in 120 s pulses under a 70 µL/min flow. Resulting sensograms were fitted to a 1:1 interaction model and kinetic constants obtained using the Biacore T200 Evaluation Software.

**Table 1.** Dissociation constants (*K*_d_ values) for the interaction of NB-AGTs with different variants of AGT as determined by SPR. In general, error propagation (from those of the kinetic analyses provided by the software) yielded errors lower than 10%, but in many cases the affinity seems to be so high that propagation is not likely reliable. Thus, estimated errors are not reported. N. Det: non-detected. Values are the average of two independent replicates.

| NB-AGT | $K_d$ (nM) | | | |
| --- | --- | --- | --- | --- |
|  | AGT-WT | AGT-LM | AGT-LM-G170R | AGT-LM-I244T |
| **1** | N. Det. | 3.8 | 20 | 0.23 |
| **2** | 0.3 | 0.3 | 0.5 | 0.2 |
| **3** | 0.5 | 0.4 | 0.5 | 0.5 |
| **4** | 0.018 | 0.02 | 0.02 | 0.024 |
| **5** | < 0.001 | < 0.001 | < 0.001 | < 0.001 |
| **6** | 10 | ~ 0.005 | 13 | 13 |

### 2.5. Hydrogen-Deuterium Exchange - Mass Spectrometry (HDX-MS)

The structural changes accompanying the binding of the six different nanobodies were followed for holo AGT WT and its variants LM and LM-G170R. AGT supplemented with a 10-fold molar excess of PLP was subjected to HDX alone or mixed 1:1 with each nanobody. PAL DHR autosampler (CTC Analytics AG, Zwingen, Switzerland) controlled by Chronos software (AxelSemrau, Sprockhövel, Germany) was used to prepare the HDX reactions. 5 µL of 10 µM protein solution (20 mM HEPES-NaOH, pH 7.4, 200 mM NaCl) were diluted into D2O-based buffer (20 mM HEPES, pD 7.4/pHread 7.0, 200 mM NaCl) at the ratio 1:9 (v/v). HDX was quenched after 20, 60, 300, 1200 and 7200 s of exchange by the addition of 50 µL cooled 1M glycine-HCl, pH 2.3. Time points 20 and 1200 s were replicated. After quenching, each sample was immediately injected into the temperature controlled LC system and digested using a nepenthesin-2 protease column (bed volume 66 µL). Digestion and desalting by 0.4% formic acid (FA) in water were driven by the 1260 Infinity II Quaternary pump (Agilent Technologies, Waldbronn, Germany) under the flow rate of 200 µL·min-1. Trapped and desalted peptides were then eluted and separated using an analytical column (Luna Omega Polar C18, 1.6 µm, 100 Å, 1.0x100 mm, Phenomenex, Torrance, CA, USA) by water-acetonitrile (ACN) gradient (10 %–45 % in 6 min; solvent A: 0.1% FA in water, solvent B: 0.1% FA, 2% water in ACN) driven by the 1290 Infinity II LC pump (Agilent Technologies, Waldbronn, Germany) under the flow rate of 40 µL·min-1. The LC-system was interfaced with an ESI source of timsTOF Pro (Bruker Daltonics, Bremen, Germany) mass spectrometer operating in the MS mode with a 1Hz data acquisition. Fully deuterated controls were used to compensate for deuterium back-exchange as described previously[40]. Acquired LC-MS data were peak picked and exported in DataAnalysis (v. 5.3, Bruker Daltonics, Bremen, Germany) and further processed by the DeutEx software [41]. Data visualization was performed using MSTools (http://peterslab.org/MSTools/index.php)[42]. For peptide identification, the same LC-MS system was used but the mass spectrometer was operated in data-dependent MS/MS mode with PASEF. The LC-MS/MS data were searched using MASCOT (v. 2.7, Matrix Science) against a customized database combining sequences of AGT WT, LM, LM.G170R, the protease and a common contaminants - cRAP.fasta (https://www.thegpm.org/crap/). Search parameters were set as follows: no-enzyme, no modifications allowed, precursor tolerance 10 ppm, fragment ion tolerance 0.05 Da, decoy search enabled, FDR ˂ 1%, IonScore ˃ 20 and peptide length ˃ 5. The HDX-MS data have been deposited to the ProteomeXchange Consortium via the PRIDE partner repository [43] with the dataset identifier PXD060227.

### 2.6. AGT enzyme activity

To determine the kinetic parameters for the transamination reaction, the purified proteins (0.2 μM) were incubated in the presence of 100 mM PLP in KP 0.1 M pH 7.4 at 25°C at increasing alanine concentrations (15–500 mM) and fixed glyoxylate (10 mM) in the absence and presence of NB-AGT-3 at two different concentrations 1 and 100 μM for 30 minutes before to start the assay. The reaction was stopped by adding TCA 10% (v/v). The pyruvate production was measured using a spectrophotometric assay coupled with lactate dehydrogenase. The pyruvate production was measured using a spectrophotometric assay coupled with lactate dehydrogenase. The Michaelis-Menten equation (equation 1) was fitted to the experimentally determine activity data (the rate constant *k*) as follows:

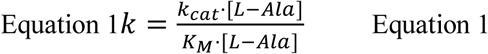

Where *k*_cat_ is the catalytic rate constant, *K*_M_ is the Michaelis-Menten constant and [L-Ala] is the actual concentration of L-Ala.

### 2.7. Cell cultures

#### 2.7.1. Cell culture, treatment and lysis

Clones of chinese hamster ovary cells stably expressing glycolate oxidase (CHO-GO) and wild-type or different variants of AGT were cultured in Ham’s F12 Glutamax medium supplemented with fetal bovine serum (10%, v/v), penicillin (100 units/ml) and streptomycin (100 mg/ml) at 37°C in a 5% CO_2_ humidified environment. The expression of AGT and GO was maintained by adding G-418 (0.8 mg/ml) and Zeocin (0.4 mg/ml), respectively, to the culture medium. To test the effects of each NBs (1C, 2C, 3C, 6C), cells were seeded in Petri dishes at the density of 9,000/cm^2^ and cultured for 7 days in the presence or in the absence of 10 µM of different compounds. At the end of treatment, cells were harvested and lysed by freeze/thawing (five cycles) in Phosphate Buffer Saline (PBS), pH 7.2 supplemented with 100 µM PLP and protease inhibitor cocktail (Complete Mini, Roche). The whole-cell extract was separated by centrifugation (13200 *g*, 10 min, 4°C) to obtain the soluble fraction. Protein concentration in the soluble fraction was measured using the Bradford protein assay.

#### 2.7.2. Western blot analyses

Five µg of soluble cell lysate was loaded on 10% SDS–PAGE, transferred on a nitrocellulose membrane and immunoblotted with anti AGT antibody (1:10.000) in 2.5% (w/v) milk in TTBS (50 mM Tris–HCl, pH 7.5, 150 mM NaCl, 0.1% Tween 20) overnight at 4°C. After three washes in TTBS, the membrane was incubated with peroxidase-conjugated antirabbit immunoglobulin G (IgG) (1:5.000) in 5% milk in TTBS for 1 h at room temperature. Tubulin (1:5000) antibody was used as loading control. Immunocomplexes were visualized by an enhanced chemiluminescence kit (ECL, Pierce Biotechnology, Rockford, IL).

#### 2.7.3. AGT activity in cell extracts

To measure the enzymatic activity in CHO cellular lysates, 30 μg of lysate were incubated with 0.5 M L-alanine and 10 mM glyoxylate at 25°C for 30 min in 100 mM KP buffer, pH 7.4 in the presence of 100 μM PLP. The reactions were stopped by adding TCA 10% (v/v). Pyruvate was determined as described in section 2.6.

### 2.8. Structural analyses

Alpha-Fold 2 was used to create structural models of all NB-AGTs [44](https://colab.research.google.com/github/sokrypton/ColabFold/blob/main/AlphaFold2.ipynb), using the procedures described recently [45]. Structural models were aligned using Pymol Multiple sequence alignment was carried out using Clustal Omega (https://www.ebi.ac.uk/jdispatcher/msa/clustalo). Accessible surface areas of the segments of AGT that were stabilized upon NB-AGT binding were displayed using the crystal structure of AGT WT (PDB code 1H0C [46]) and calculated using GetArea (https://curie.utmb.edu/getarea.html) [47].

## 3. Results

### 3.1. Generation of Nanobodies towards AGT (NB-AGTs)

We successfully generated nanobodies against AGT-LM (named as NB-AGT along the manuscript) which also bind to AGT-WT based on ELISA measurements. Sequence alignment of the six NB-AGT allowed to determine the location of hypervariable loops (CDR1-3, Figure 1) responsible for binding of NB to its target [20]. According to structural predictions by AlphaFold 2 [44], all NB-AGT fold similarly into a structure rich in β-sheet with the largest structural divergence between them in the CDR3 (Figure 1). All NB-AGT were produced in *E.coli* and purified (Figure S1), and seem to be well folded, with hydrodynamic radius in the 1.9-2.1 nm (Figure S2A), a rich content in β-sheet based on the far-UV CD spectra (Figure S2B) and showing high thermostability and cooperative denaturation (Figure S2C). Analyses of thermal denaturation of NB-AGTs using a pseudo two state reversible unfolding showed that a two-state unfolding model described well their thermal unfolding (Figure S3 and Table S1), as previously reported for NB-AGT-1, NB-AGT-6 and NB-AGT-2 wild-type sequences and several mutants of NB-AGT-2 [39,45]. Different thermostability of the NB-AGTs likely reflect different thermodynamic stabilities at lower temperatures (25°C or 37°C) as recently proposed [39,45]. The expected change in enthalpy for reversible denaturation of proteins with the size of NB-AGT lies within 80-90 kcal·mol^-1^ based on structure-energetic relationships [48] which agrees well with experimental enthalpies (average of 76 ± 12 kcal·mol^-1^ for the six NB-AGTs, Table S1), thus supporting that all NB-AGTs are natively folded and undergo extensive denaturation upon thermal unfolding.

### 3.2. Tight Binding of NB-AGT to pathogenic and non-pathogenic AGT variants

We then evaluated the affinity of NB-AGT for different AGT variants by two different approaches. We first determined the effect of NB-AGTs on the stability of AGT-LM (Figure 2 and S4). All NB-AGT affected the thermal stability of AGT-LM, with the mildest effect in NB-AGT-2. In all cases (except for NB-AGT-2), the maximal effect was found around a 1:1 ratio of NB-AGT and AGT-LM, supporting a tight binding (Figure 2). We must note that the finding of two NB-AGTs that destabilized AGT-LM *in vitro* against thermal denaturation implies interaction with but not necessarily destabilization of AGT-LM under physiological conditions (the NB-AGT may change the kinetic route of denaturation at high temperatures; note that these occurred over 60°C and these effects might be quite different at room or physiological temperature).

**Figure 2.**
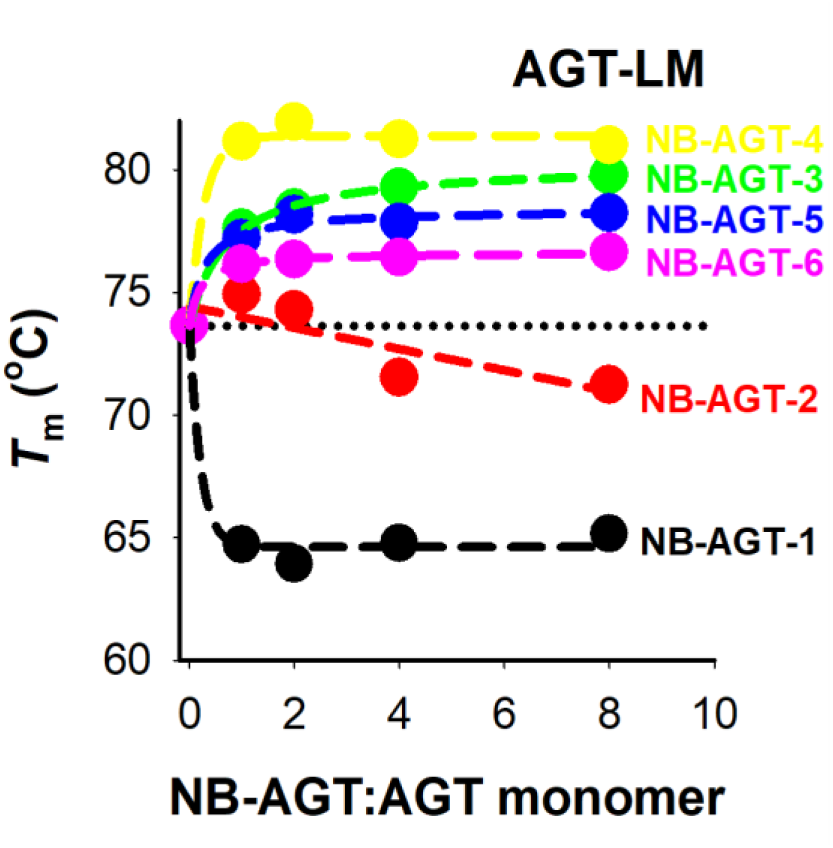
DSC analyses confirm the interaction of NB-AGTs with AGT-LM. Scans were run from 5 to 130 °C at a scan rate of 2 °C·min^-1^. The experiments were carried out in 20 mM HEPES-NaOH, 200 mM NaCl pH 7.4. Protein concentration was 5-40 µM for the NB-AGTs and 5 µM (in monomer) for the AGT in the presence of 25 µM of pyridoxal 5′-phosphate (PLP). The dotted horizontal line represents the *T*_m_ of AGT-LM in the absence of NB-AGT. Colored solid lines were hyperbolic fittings to experimental data (colored symbols, *T*_m_ at different NB-AGT:AGT ratios) with no strict theoretical meaning.

To quantitatively estimate the binding affinity of NB-AGTs for different variants of AGT (the WT, the minor allele LM containing the P11L and I340M variations, and the minor allele containing the two most prevalent mutations causing PH1, [4,27,31], we used surface plasmon resonance (SPR) (Figure 3, S5 and Table 1). In general, all NB-AGT bound tightly to the different variants, with *K*_d_ values ranging from nM to even below pM (Table 1). Interestingly, some NB-AGT seemed to bind with different affinities to specific AGT variants (e.g. NB-AGT-1 and 6, see Table 1). Therefore, NB-AGTs bind very tightly to all AGT variants (with the only exception of NB-AGT-1 and AGT-WT) and in some cases the affinity depends on the AGT variant tested.

**Figure 3.**
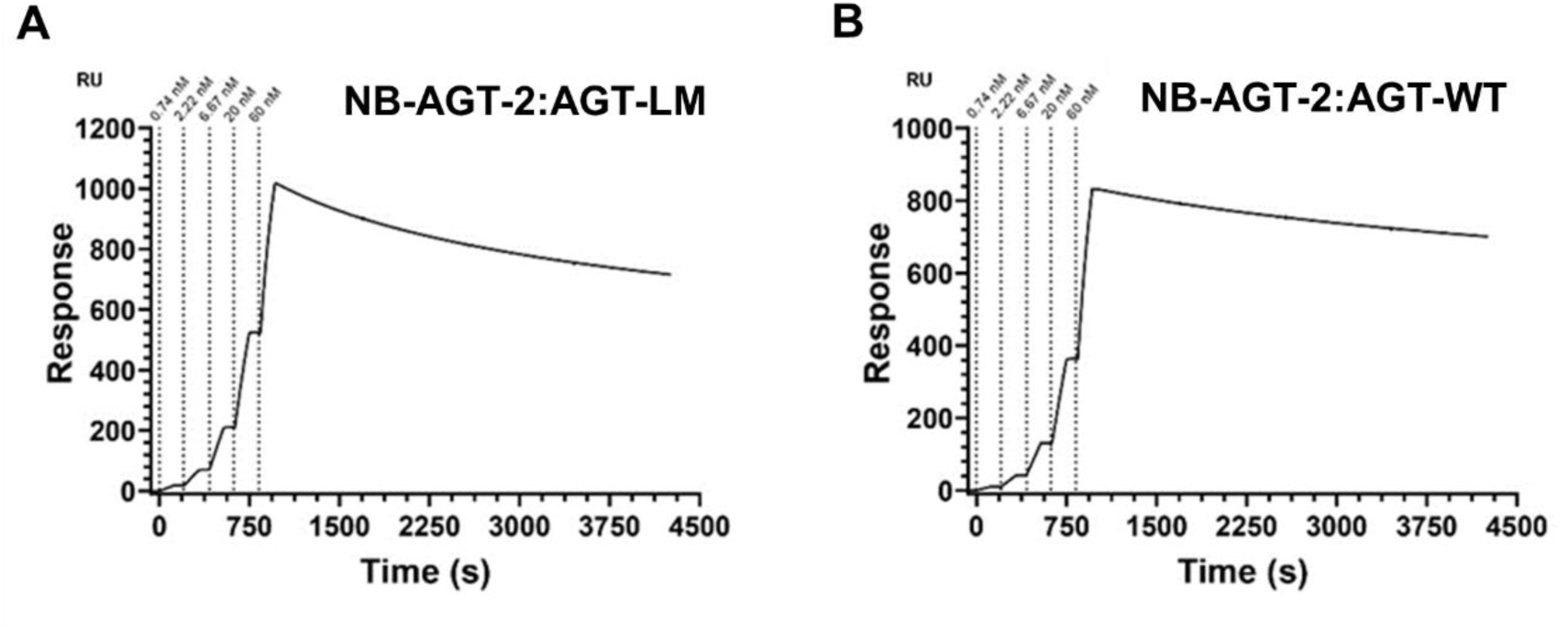
The interaction of NB-AGT-2 with AGT-LM andAGT-WT analysed by SPR. Representative sensograms corresponding to a Single Cycle Kinetics experiment performed with NB-AGT-2 as substrate and AGT-LM (A) and AGT-WT (B) as analytes. Five analyte concentrations were sequentially injected during 120 s with 30 s stabilization. Transients were monitored for 3300 s. Data ware analysed using Biacore T200 Evaluation Software.

### 3.3 Structural basis of the NB-AGT interaction with AGT variants

To provide structural insight into the interaction of holo-AGT variants (WT, LM and LM-G170R) with the NB-AGT (1 to 6), we have carried out HDX-MS studies with the AGT variants in the presence or absence of each NB-AGT. HDX data were derived from 252 (WT), 249 (LM) and 250 (LM-G170R) peptides, covering 98.5, 97.5 and 97.2% sequence, respectively. The average peptide length was between 10.2 and 10.4 and average redundancy was 6.6-6.7 (Figure S6). The effect of NB-AGT binding on the kinetics of deuterium incorporation (%D vs. time) for all these experiments can be found in Figure S7.

We compared the HDX of a given AGT in the absence/presence of the NB-AGT, to detect those regions in AGT that changed local thermodynamic stability as the Δ%D_av_ values (cut-off ≥ |10|) [17,49,50]. We may refer to these regions in AGT for a given NB-AGT as the NB *binding sites*, even though regions far from the interface between the NB and AGT might be also stabilized upon binding [17]. This approach provided a consistent picture of the interaction of AGT with the NB-AGTs, and these interactions were observed in all cases, except for AGT-WT and NB-AGT-1 (Figure S8 and Table S2), were consistent with our SPR analyses (Table 1). Although some small differences were found for the *binding site* of a given NB-AGT and AGT-WT, LM and LM-G170R (Figure S8 and Table S2), a consensus interaction region in AGT could be determined considering the experimental uncertainty for each NB-AGT (Figure 4). When these interacting regions were analyzed, we may classify the *binding sites* for the six NB-AGT in four groups (Figure 4): 1) NB-AGT-1 binding decreases the stability of five segments in the NTD, that could be associated with the strong thermal destabilization exerted by this NB-AGT on AGT-LM. The average solvent accessible surface area of this binding site is of ∼13% using GetArea [47] and the AGT structure with the PDB code 1H0C [46]; 2) NB-AGT-2 binding stabilizes a small region in the NTD of AGT (residues 173-178) and this site is more solvent-exposed (average of ∼34% of ASA); 3) NB-AGT-3 to 5 stabilized the same three regions in the NTD (average of ∼32% of ASA), whereas NB-AGT-3 and 4 also stabilized a segment in the CTD (average of ∼23% of ASA); 4) NB-AGT stabilize three solvent-exposed regions in the CTD (average of ∼35% of ASA). The stabilization of the region 303-309 is remarkably large, with Δ%D_av_ values over 50 (Figure S7-S8).

**Figure 4.**
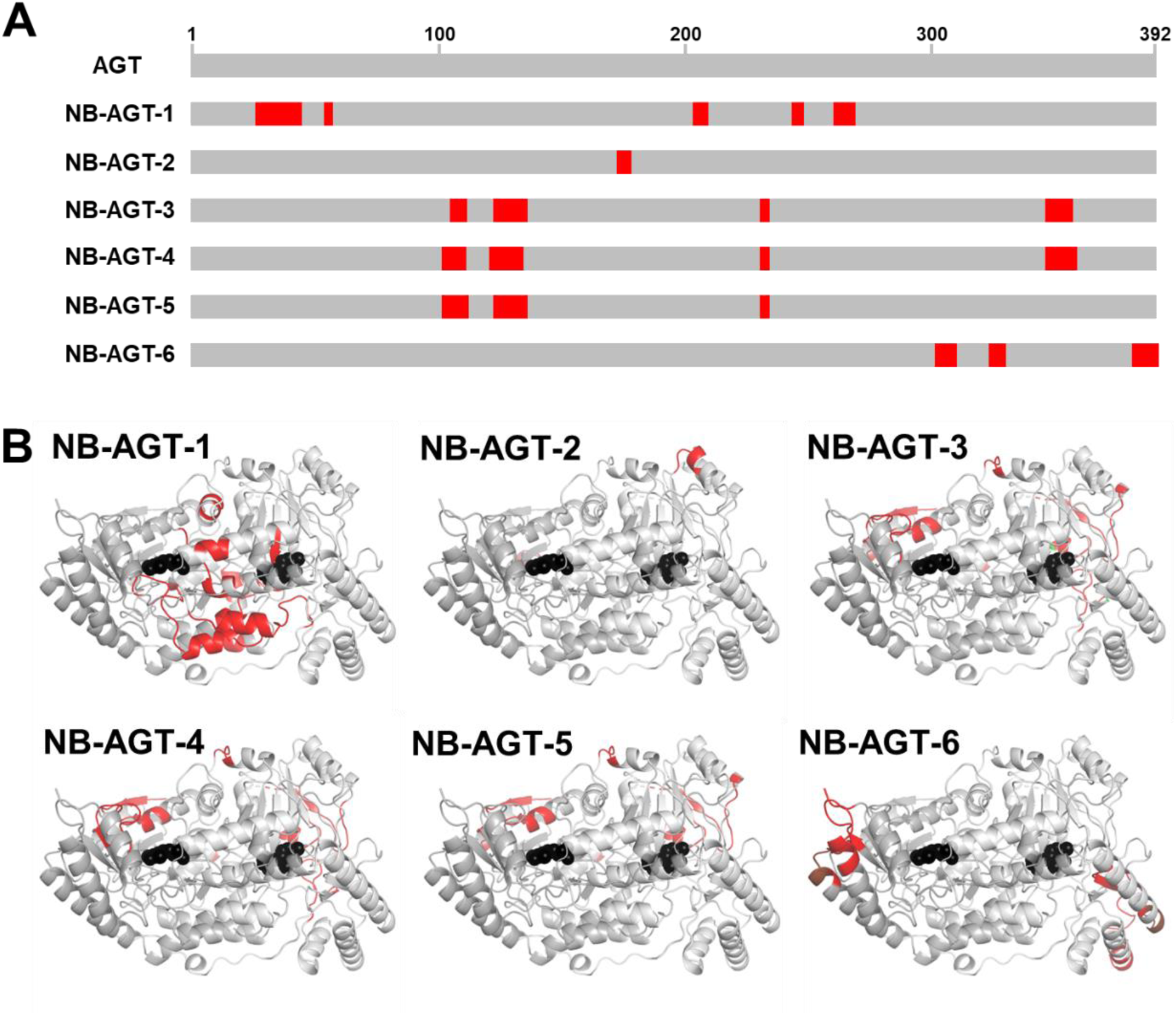
Consensus NB-AGT *binding sites* in AGT variants as determined by HDX-MS. A) Regions whose stabilities are affected by NB-AGTs binding represented in the AGT sequence. These consensus regions were determined from analyses of AGT WT, LM and LM G170R variants (Table S2). B) Structural location of the putative *binding sites* (in red). PLP is shown as black spheres. The structure used for display has the PDB code 1H0C [46].

### 3.4. NB-AGT-3 increases the specific activity of AGT-LM-G170R and LM-I244T in transfected CHO cells

To test the potential of NB-AGTs to correct AGT misfolding (aggregation), we investigated their effects on AGT protein levels (Figure 5) and activity (Figure 6) in transfected CHO cells. NB-AGTs were added to the cell cultures as purified proteins with a potent cell penetrating peptide [37] at the N-terminus of the NB-AGT sequence and containing the canonical PEX5P peroxisomal targeting sequence (-SKL) at their C-terminus [51]. We named these modified versions of the as CPP-NB-AGTs. Two CPP-NB-AGTs concentrations were tested (1 and 10 µM), using four different CPP-NB-AGT variants with different binding modes based on our HDX-MS analyses (Figure 4). Two non-pathogenic AGT variants (WT and LM) and the two most common PH1-causing mutants (LM-G170R and LM-I244T) were studied. AGT protein levels and activities were determined after a week of incubation with NB-AGTs. Regarding AGT protein levels, the exposure of transfected CHO cells to the CPP-NB-AGTs did not significantly enhance protein levels (in fact, in some cases seemed to modestly reduce them; Figure 5). In the case of AGT activity, we only observed a modest increase in activity for the AGT variants LM-G170R and LM-I244T when experiments at 1 and 10 µM CPP-NB-AGT-3 were compared (Figure 6; not statistically significant, p > 0.1 using a two-tailed *t*-test). Notably, the specific activity of LM-G170R and LM-I244T was increased in the presence of 10 µM CPP-NB-AGT-3 (to 143 ± 12 % and 139 ± 28 % vs. the same variant without NB-AGT-3). These results suggest that the treatment with NB-AGT-3 increased the fraction of functional AGT. To test for a possible inhibitory effect of CPP-NB-AGTs on AGT activity, we compared the steady-state kinetics of WT and LM-G170R variants in the absence or the presence of up to 100 µM of CPP-NB-AGT-3 (Figure S9). No significant inhibitory effects were observed (Table S3).

**Figure 5.**
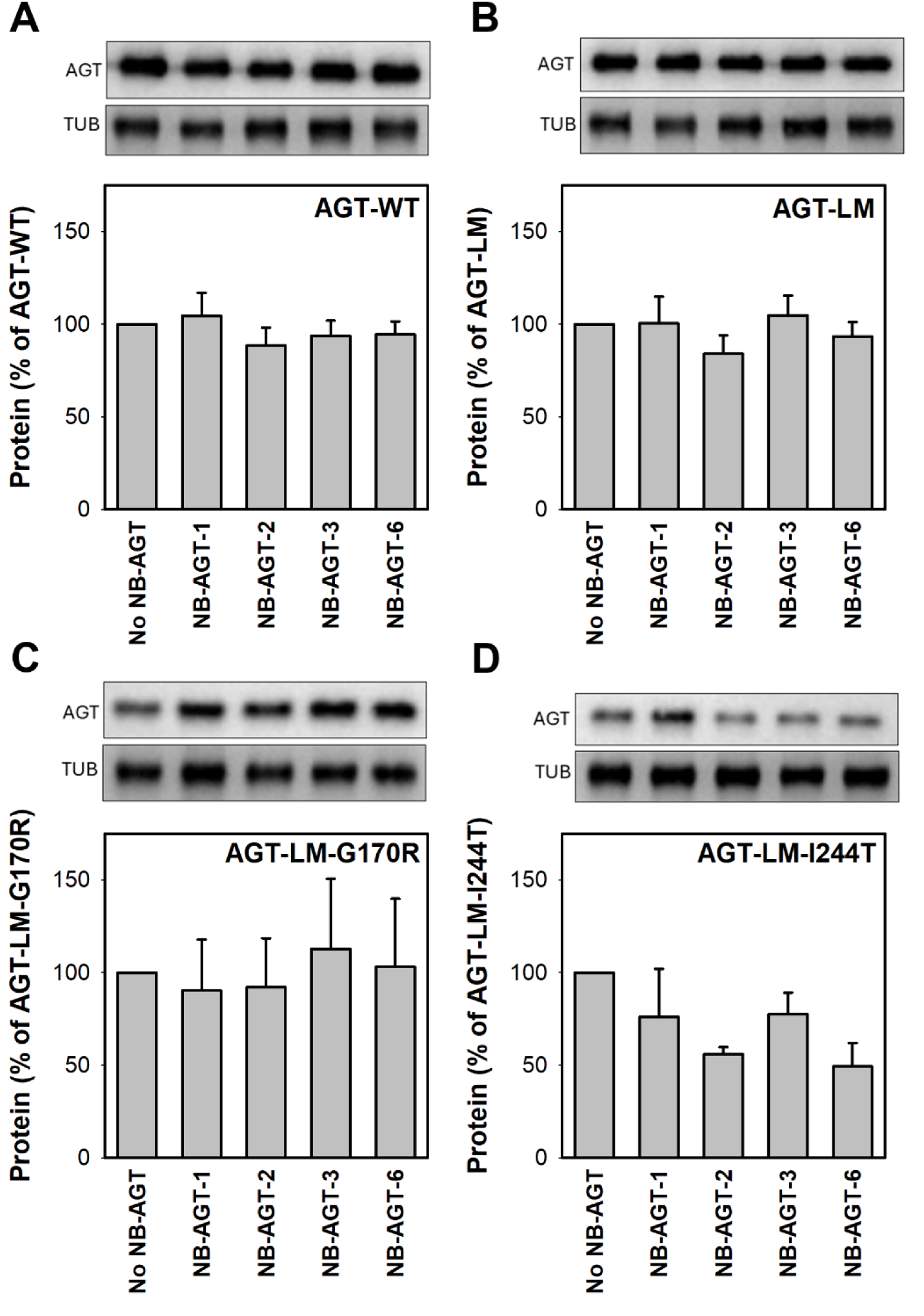
The effect of CPP-NB-AGT on the protein levels in transfected CHO cells. Cells were incubated with each NB-AGT at 10 µM in the cell medium for one week. The AGT and tubulin (TUB) protein levels were determined in soluble extracts by western-blot (upper insets) whereas normalized levels (using AGT and TUB) of AGT protein were normalized in each panel using the levels of the corresponding variant without NB-AGT (lower insets; data are mean±s.d. from 3-4 independent experiments). Panels show data for AGT-WT (A), AGT-LM (B), AGT-LM-G170R (C) and AGT-LM-I244T (D).

**Figure 6.**
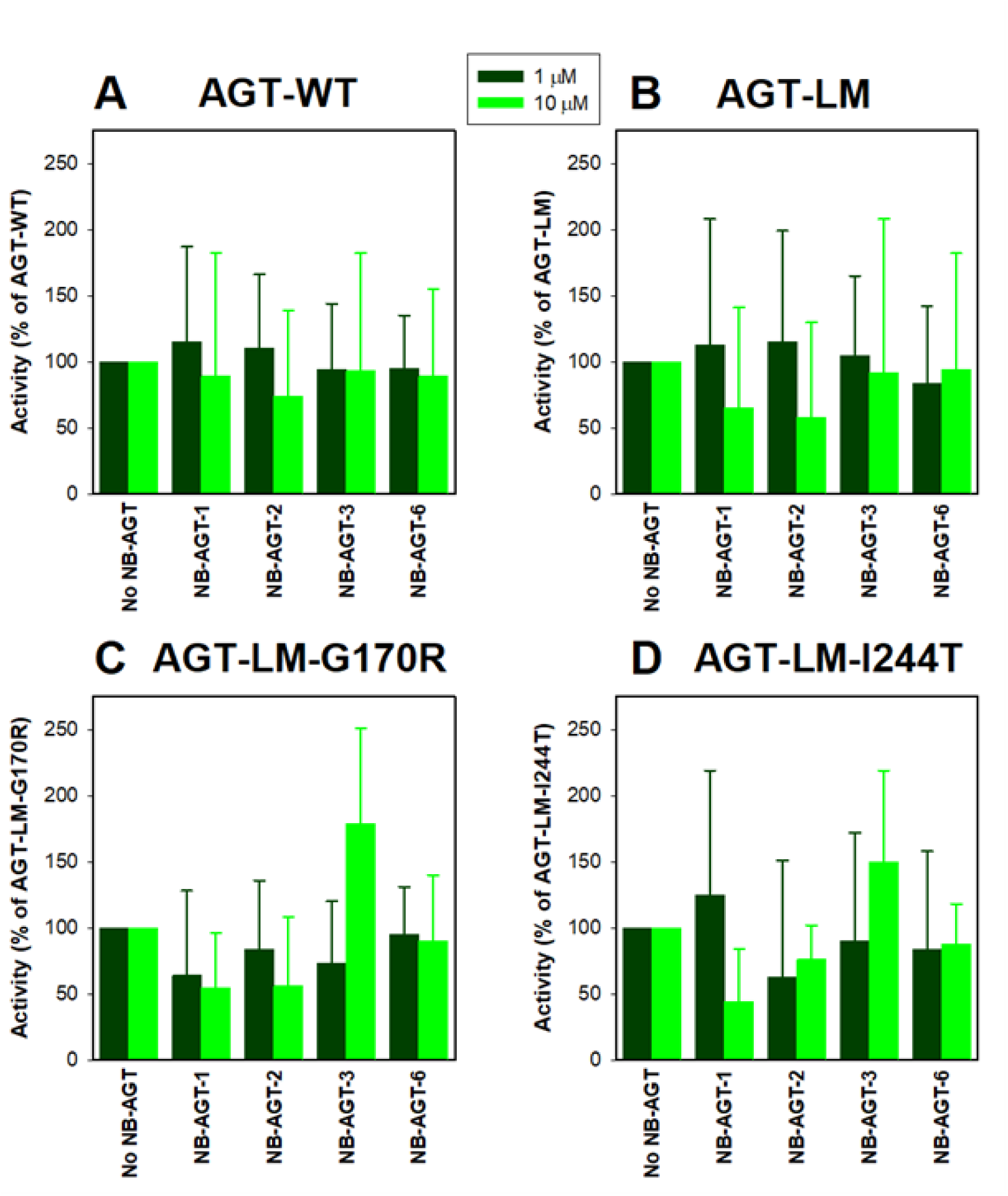
The effect of CPP-NB-AGT on the AGT activity in transfected CHO cells. Cells were incubated with each NB-AGT at 1 (dark green) or 10 (light green) µM in the cell medium for one week. The AGT activities were determined in soluble extracts by western-blot and normalized using the corresponding variant without NB-AGT. Data are mean±s.d. from three independent experiments). Panels show data for AGT-WT (A), AGT-LM (B), AGT-LM-G170R (C) and AGT-LM-I244T (D).

## 4. Conclusions

Caplacizumab (commercialized by Sanofi) was the first NB approved by the FDA and EMA for the treatment of a human disease (acquired thrombotic thrombocytopenic purpura) [52–54]. A wide variety of NB are under development to prevent protein:protein interactions associated with disease [20,52,55,56]. In this work, we provide the first evidence of a NB (NB-AGT-3) that corrects a conformational disease with loss-of-function (in the case of PH1, the two most common disease-causing mutants that enhance AGT aggregation). To potentially apply NB-AGTs to treat PH1, multivalent NBs could be generated with several copies of a single nanobody increasing its potency [25,54], and humanized versions obtained by relatively simple engineering procedures [57].

Considering the ADME guidelines [58], A(bsorption) and D(istribution) were successfully targeted in our work. First, nanobodies are easy to be produced, are extremely stable at room temperature [20,35,39,45,59] so storage and delivery issues as therapeutics may outweight conventional antibody-based approaches [20,36,39,52,55]. Second, NB-AGTs largely outperform other small-molecule based approaches in terms of binding affinity for the native state (by at least 4 orders of magnitude) [14,60,61]. From a basic mass-action principle, the stabilization of the native state would be in principle greater [62] using NBs than those observed for small molecules. Even though AGT1 folding and misfolding are known to be complex [27–29,32,61,63,64], our results showed that AGT aggregation and particularly misfolding, can be corrected by nanobodies. We plan to modify our approach to target CPP-AGT-NB-AGT to target hepatocytes specifically [65–67] using mouse models of PH1 available in our laboratories [68,69].

## Abbreviations

AGT: alanine:glyoxylate aminotransferase 1
AGT-LM: AGT minor allele
AGT-WT: AGT major allele or wild-type
BB: binding buffer
CD: circular dichroism
CPP: cell-penetrating peptide
CPP-NB-AGT: NB-AGT with a CPP attached
DLS: dynamic light scattering
DSC: differential scanning calorimetry
ELISA: enzyme linked immusorbed assay
HEPES: 2-[4-(2-hydroxyethyl)piperazin-1-yl]ethanesulfonic acid
IMAC: immobilized metal affinity chromatography
K_d_: dissociation constant
LBK: Luria-Bertani medium with kanamycin
NB: nanobody
NB-AGT: nanobody raised against AGT
NTA: nitriloacetic acid
PH1: primary hyperoxaluria type 1
PLP: pyridoxal 5′-phosphate
RU: response unit
SDS-PAGE: gel electrophoresis in the presence of sodium dodecylsulphate
SPR: surface plasmon resonance

## Acknowledgements

A.L.P. acknowledges funding from Consejería de Economía, Conocimiento, Empresas y Universidad, Junta de Andalucía (Grant number P18-RT-2413).P.M. acknowledges support from the MEYS project OP JAK INTER-MICRO (CZ.02.01.01/00/22_008/0004597). Access to the Instruct-CZ center (BioCeV), was supported by CIISB (LM2023042 and CZ.02.01.01/00/23_015/0008175) and Instruct-Internship to J.L.P-G. (PID 2545). Mario Cano-Muñoz was supported by Juan de la Cierva Postdoctoral Research Program from the Spanish Research Agency Grant: JDC2022-049681-I funded by MICIU/AEI/10.13039/501100011033.

## Author contributions

Conceptualization: A.L.P.; Data curation: M.D., P.M., B.C. F.C-L. and A.L.P.; Formal analysis: M.D., P.M., B.C., F.C-L. and A.L.P.; Funding acquisition: E.S., B.C. and A.L.P.; Investigation: A.G-M., M.D., J.L.P-G, S.G., M.C-M, P.V., D.S.L.,; Methodology: M.D., M.C-M., P.M., F.C-L, B.C. and A.L.P.; Project administration: A.L.P.; Resources: M.C-M., E.S., P.M., B.C. and A.L.P.; Software: M.C-M., P.M. and A.L.P.; Supervision: A.L.P.; Validation: P.M., B.C. and A.L.P.; Visualization: A.L.P; Writing – original draft: A.L.P; Writing – review and editing: M.D., P.M., E.S., B.C. and A.L.P.

