## Supplementary Data for "Nanobodies as therapies for loss-of-function misfolding diseases"

### Supplementary Information

### Methods

#### Far-UV circular dichroism (CD) spectroscopy

CD spectra were registered in a Jasco J-710 spectropolarimeter (Tokyo, Japan) with a Peltier element for temperature control. Before the acquisition of far-UV CD spectra, protein samples were buffer exchanged to 20 mM K-phosphate pH 7.4 using PD-10 columns. Spectra in the far-UV range were acquired using 40 µM of NB-AGTs in a 1 mm path-length quartz cuvette. These spectra were registered in the 210-260 nm range, at 25 ºC with a scan rate of 100 nm·min$^{-1}$. Five spectra of each sample were acquired and averaged, and the corresponding blank (containing the same buffer but no protein) was acquired under the same conditions and subtracted from those of protein samples. To calculate mean molar residue ellipticities $[\Theta]_{MRW}$ we used the equation S1:

$$[\Theta]_{MRW} = \frac{MRW \cdot \Theta}{10 \cdot l \cdot c} \qquad \text{Equation S1}$$

Where MRW is the molecular weight of each nanobody divided by the number of residues minus 1, Θ is the observed ellipticity of the sample in mdeg, l is the pathlength of the cuvette in cm and c the protein concentration in mg·mL$^{-1}$.

#### Dynamic light scattering (DLS)

DLS was carried out in a DynaPro MSX instrument (Wyatt, Santa Barbara, CA, USA) using 1.5 mm path length cuvettes and 15 µM NB-AGTs in 20 mM K-phosphate pH 7.4 at 25 ºC. 30 measurements with an acquisition time of 10 s were acquired for each nanobody and used to determine the hydrodynamic radius assuming spherical scattering particles (Stokes-Einstein approach).

**Figure S1. Purity and integrity of NB-AGTs. A)** 15% SDS-PAGE gel of samples obtained from IMAC+SEC for all six NB-AGTs. B) HPLC/ESI-MS analyses showing high purity and molecular weight consistent with each NB-AGT sequence.

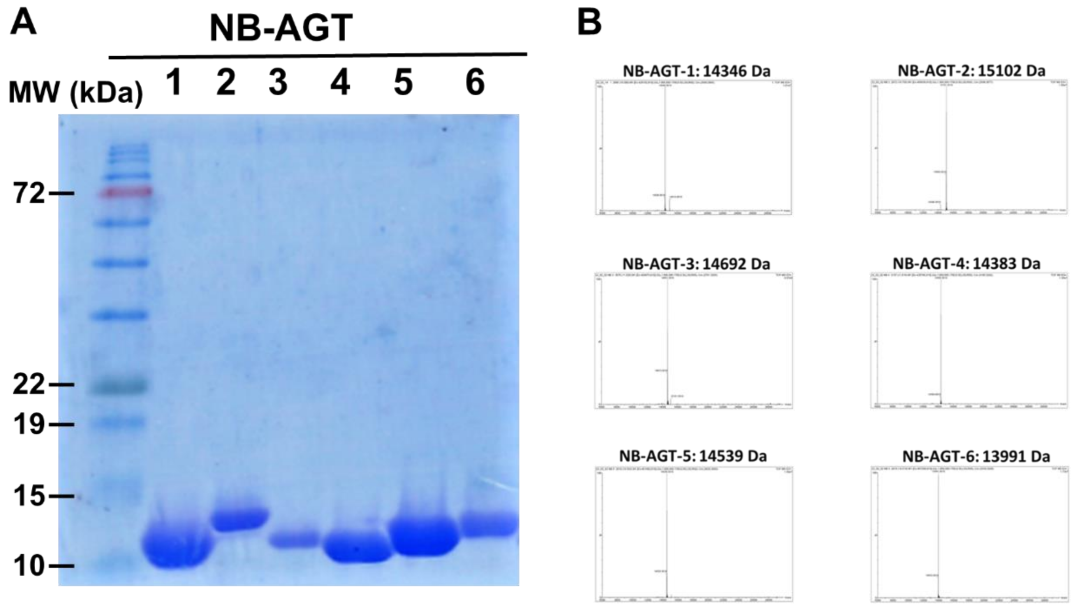


**Figure S2. Biophysical studies analyses support a well-folded conformation for the 6 NB-AGTs.** A) DLS studies of size populations for NB-AGTs. B) far-UV CD spectra of NB-AGTs. C) Thermal denaturation analyses by DSC showing that NB-AGT are well folded and highly stable proteins.

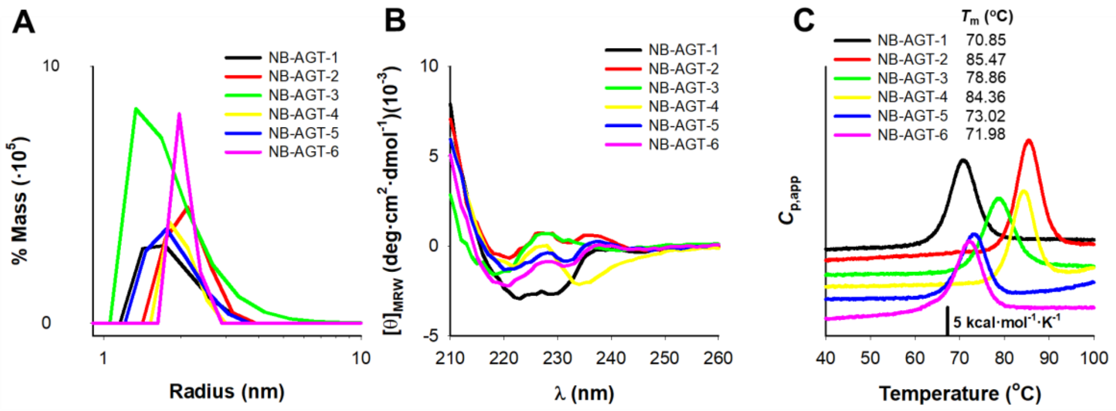

**Figure S3. DSC thermograms for the six NB-AGTs analyzed using a reversible pseudo two-state unfolding model.**

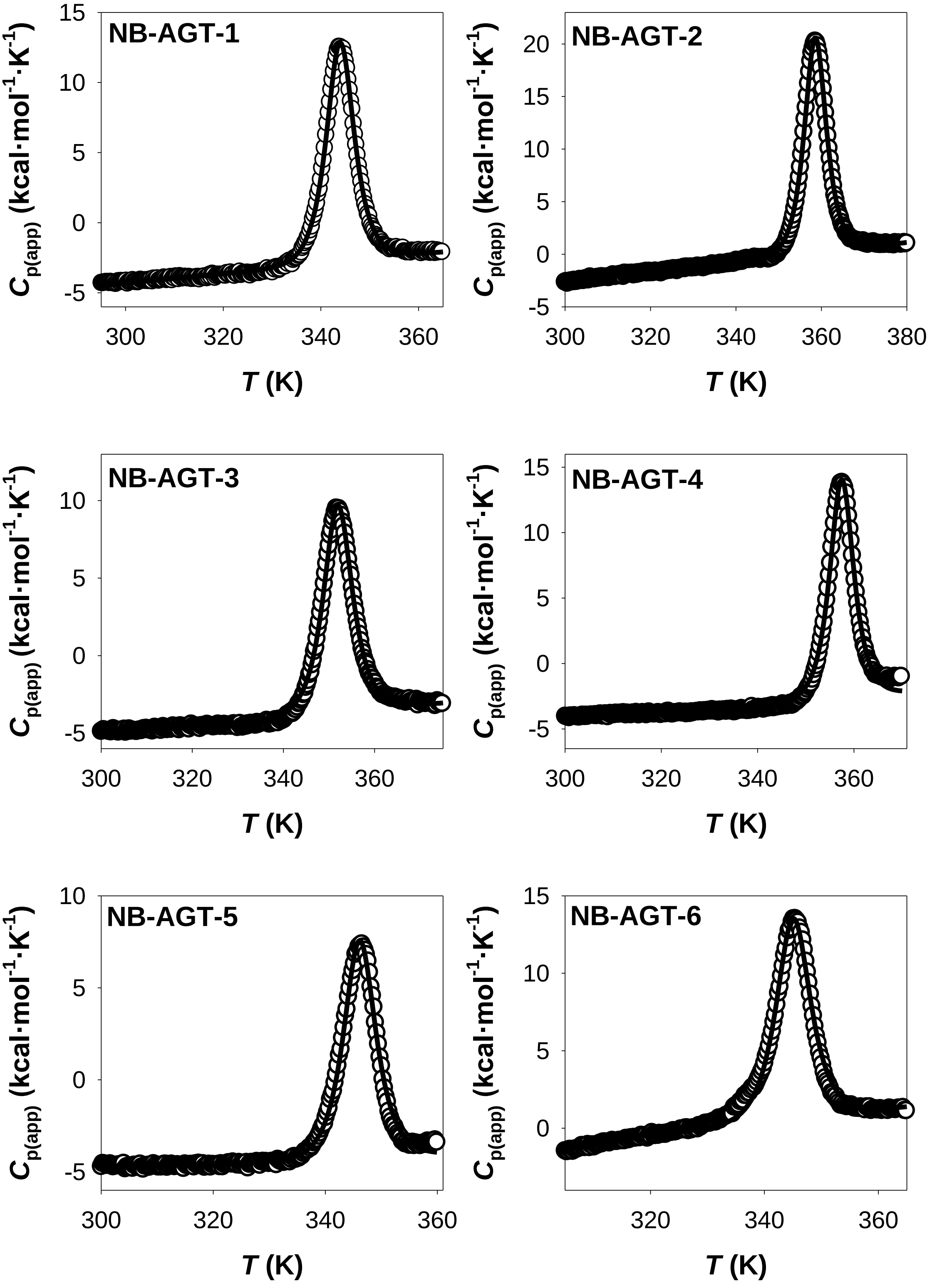

**Figure S4. DSC thermograms for AGT-LM and NB-AGTs alone and in complex mixed with two different ratios.** Insets show the differential plots between AGT-LM alone and in the presence of NB-AGTs.

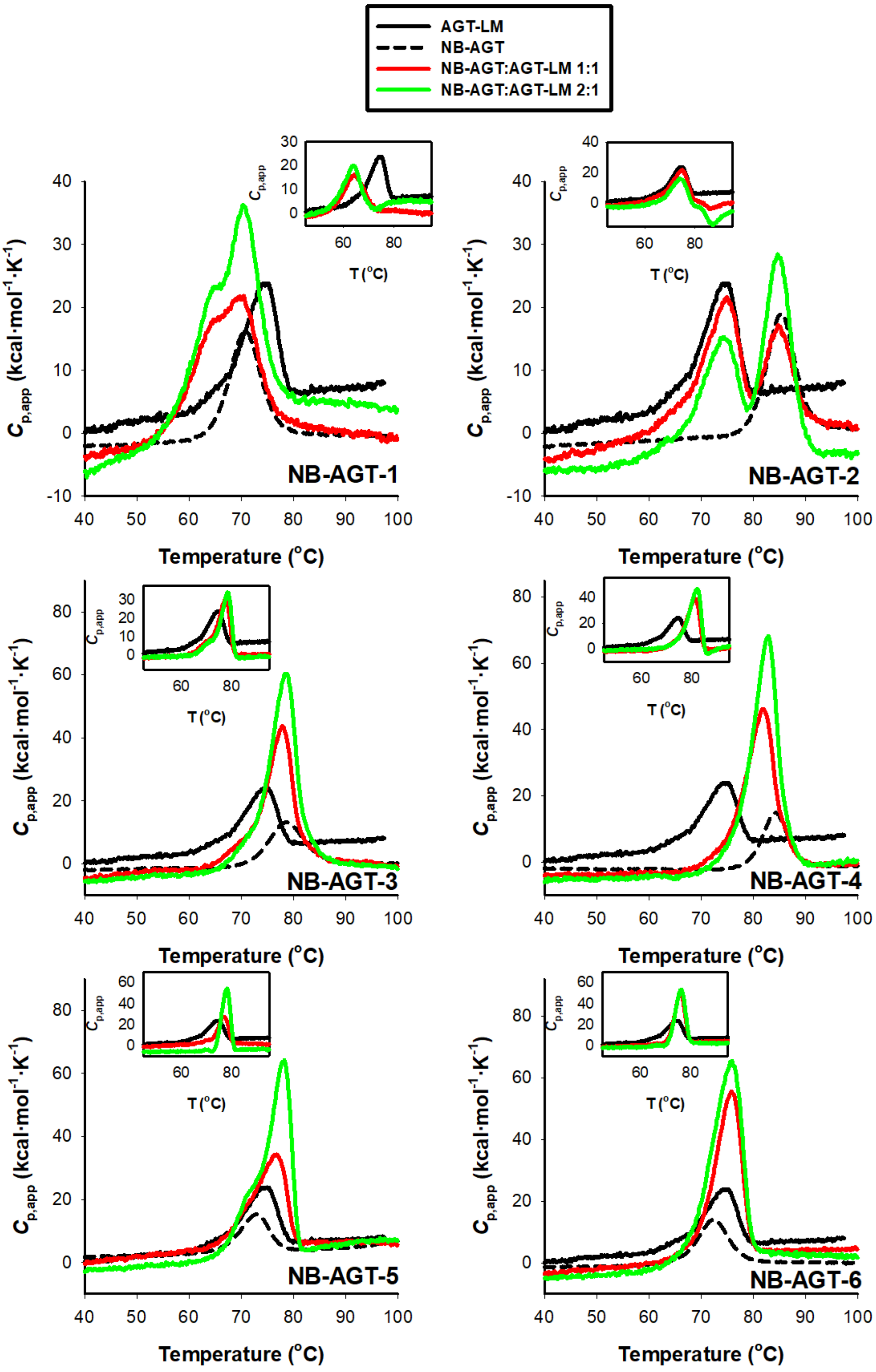

**Figure S5. Reproducibility of SPR analyses.** Blank subtracted SPR sensograms corresponding to a Single Cycle Kinetics experiment performed with different NB-AGTs immobilized in the chip and AGT-WT as analyte. Five analyte concentrations were sequentially injected during 120 s with 30 s stabilization. Transients were monitored for 3300 seconds. Data were analysed using Biacore T200 Evaluation Software. The kinetics were analysed using several cycles: two start-up cycles, preparatory to condition the chip, and a sample cycle followed by a blank one, duplicated. Five analyte concentrations were used, injected during 120 s with 30 s stabilization, without regeneration in between, ranging from 0.74-60 nM.

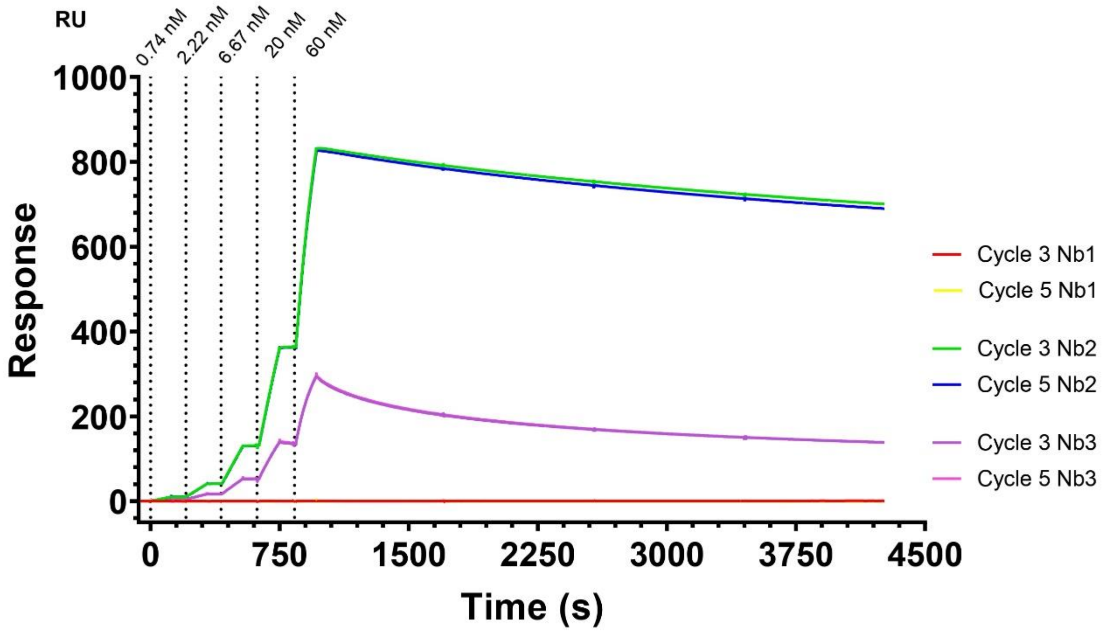

**Figure S6. Sequence coverage of Holo-AGT WT, LM and LM-G170R.** Peptides providing HDX-MS data plotted against the AGT sequence, representing nearly complete sequence coverage (97-99%).The secondary structure elements are shown above the sequence. Blue bars are the individual peptides for WT, red for LM and green for LM-G170R.

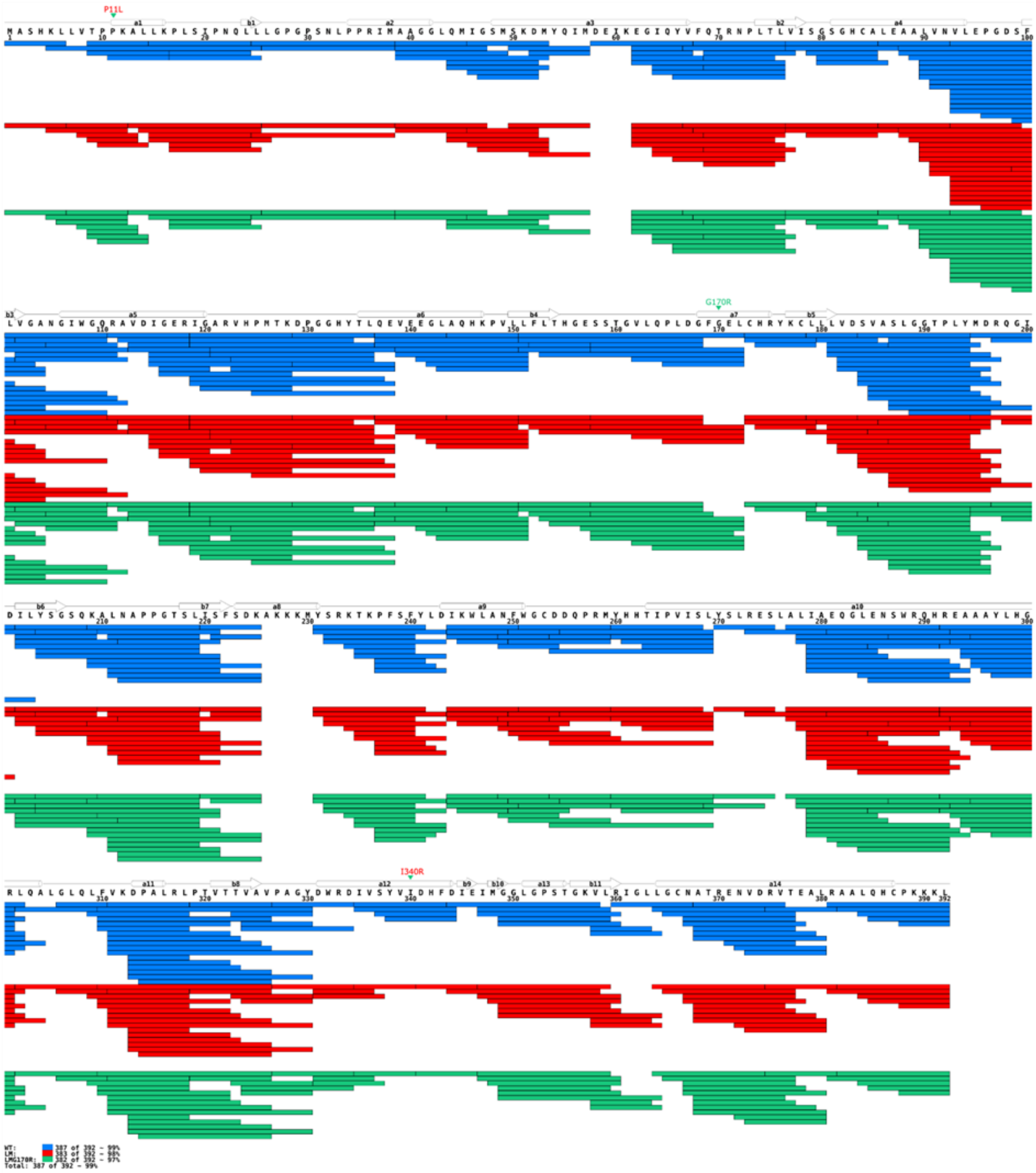

**Figure S7. Structural changes in AGT WT, LM and LM-G170R induced by nanobody binding.** Differences in deuteration between the free AGT and AGT bound to one of the six tested nanobodies shown as heat maps, where red means higher deuteration (deprotection) after nanobody binding and blue protection upon the interaction with nanobody. Maps are shown for WT (A), LM (B) and LM-G170R (C). Locations of the three mutations are shown in grey above each heat map. Areas not covered by the HDX data are shown in grey.

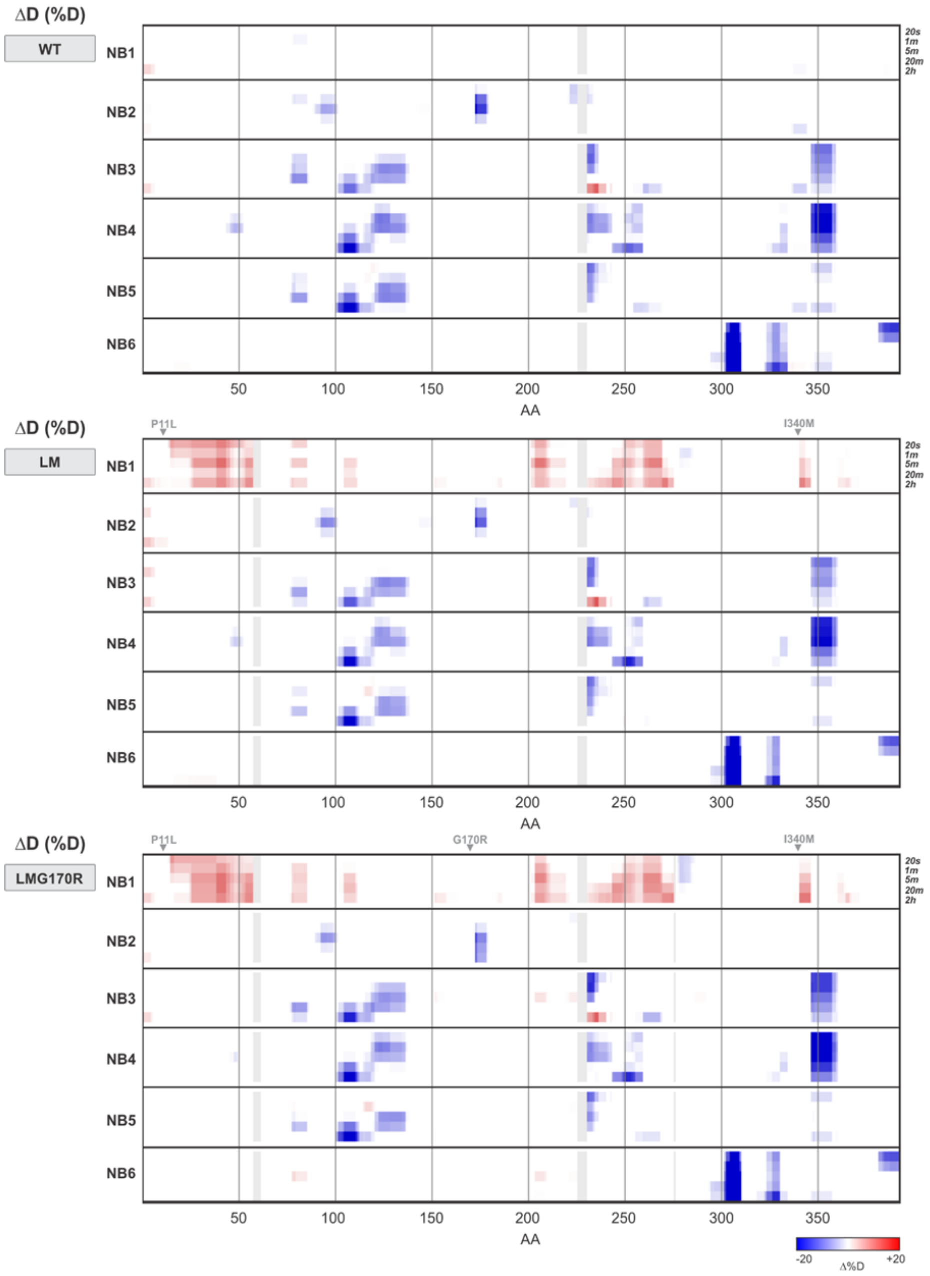

**Figure S8. Consensus AGT regions (*binding sites*) with changes in stability ($\Delta\%D_{av} \geq |10|$) upon NB-AGT binding.** Each panel shows a consensus AGT region upon interaction with a NB-AGT, and the $\Delta\%D_{av}$ values for each consensus region is shown for individual AGT variants.

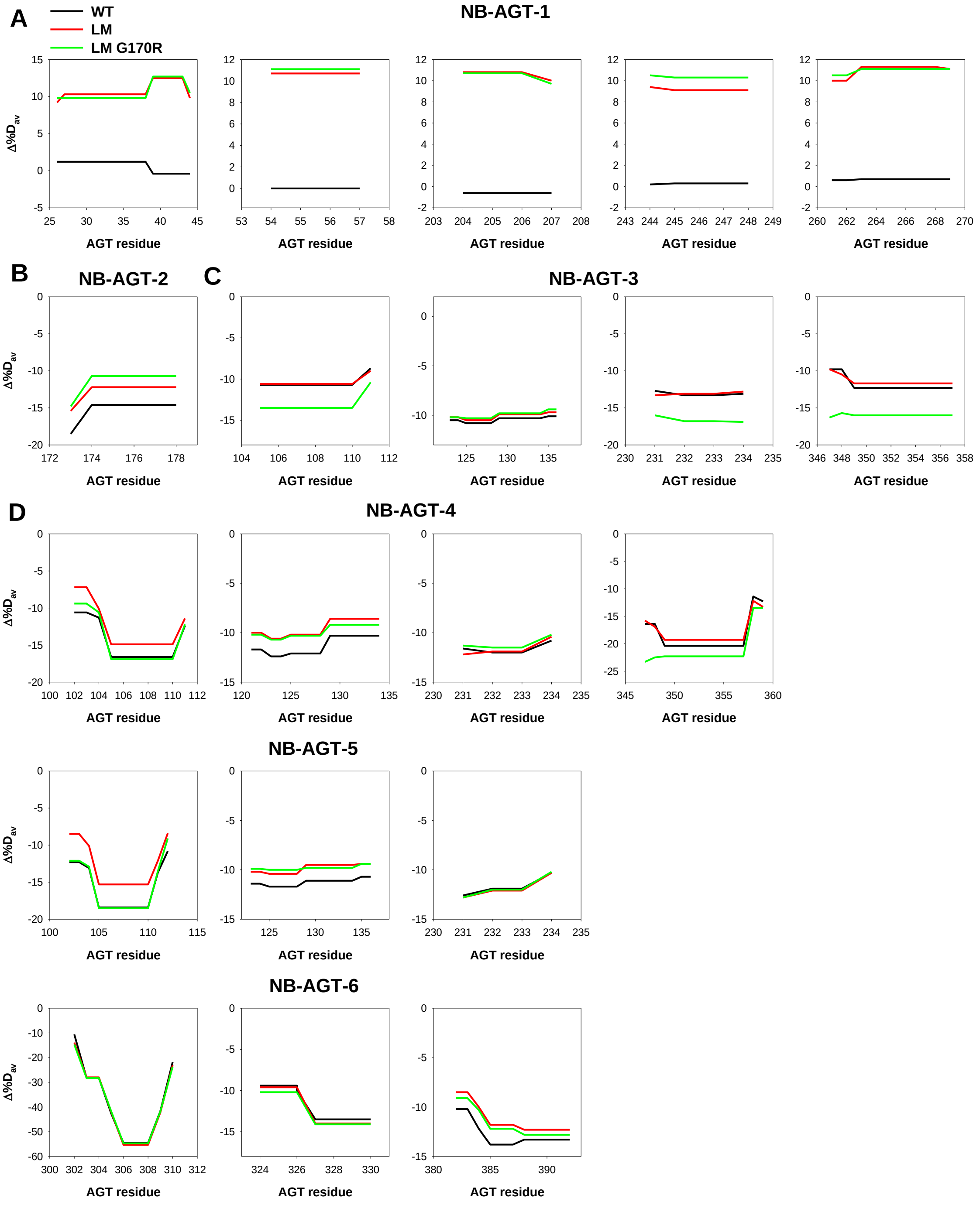

**Figure S9. Michaelis-Menten plots of AGT WT (A) and LM-G170R (B) activity in the absence and presence of the Nanobody 3 at two different concentrations. A) and B)** Values of initial velocity (nmol product/time/nmol enzyme) as a function of L-alanine concentration (15-500 mM) in the presence of 10 mM glyoxylate. The enzyme concentration was 200 nM and assays were performed in the presence of 100 μM exogenous PLP in potassium phosphate 0.1 M pH 7.4 at 25 °C.

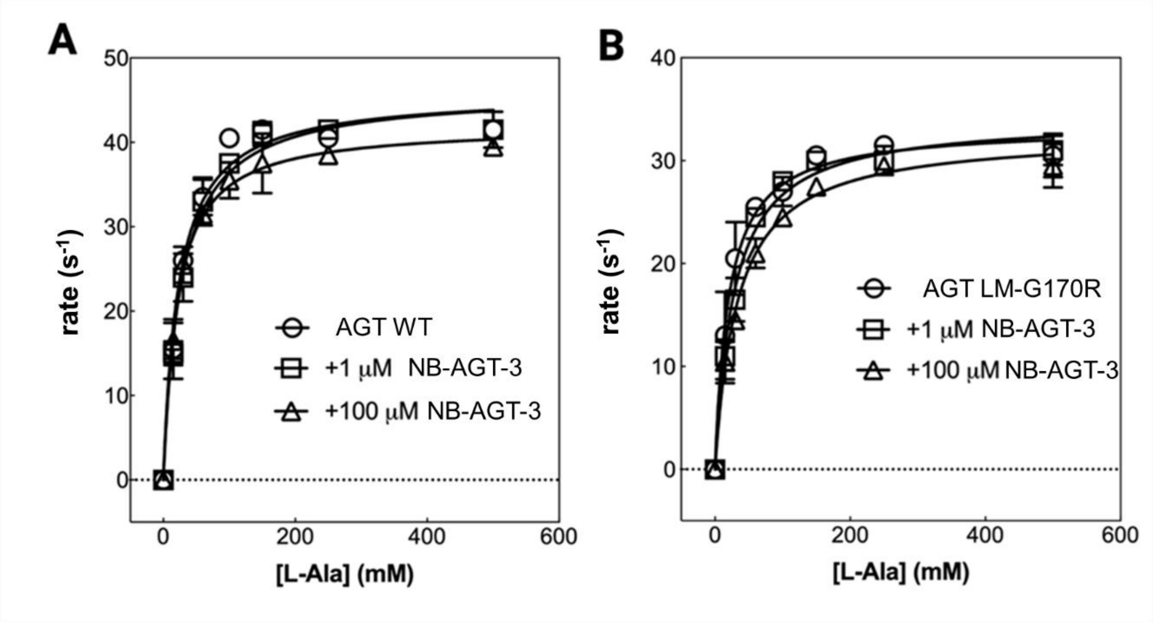

**Table S1. Thermodynamic denaturation parameters for NB-AGTs using a pseudo two-state unfolding model.** An average value of $\Delta H_{cal}$ / $\Delta H_{VH}$ of 0.8 ± 0.1 was obtained for this set of NB-AGTs, supporting that a reversible two-state unfolding model describes well their thermal denaturation (this value should be 1 for an ideal two-state model).

| NB-AGT | Thermodynamic parameters | | | |
|---|---|---|---|---|
| | $T_m$ (K) | $\Delta H_{cal}$ (kcal·mol$^{-1}$) | $\Delta H_{VH}$ (kcal·mol$^{-1}$) | $\Delta H_{cal}$ / $\Delta H_{VH}$ |
| **1** | 344.80 ± 0.01 | 78.7 ± 0.2 | 93.6 ± 0.2 | 0.84 ± 0.01 |
| **2** | 358.62 ± 0.01 | 92.7 ± 0.1 | 113.0 ± 0.2 | 0.82 ± 0.01 |
| **3** | 351.99 ± 0.01 | 75.2 ± 0.2 | 87.7 ± 0.3 | 0.86 ± 0.01 |
| **4** | 357.51 ± 0.01 | 73.2 ± 0.3 | 115.6 ± 0.6 | 0.63 ± 0.01 |
| **5** | 346.17 ± 0.01 | 59.7 ± 0.3 | 94.1 ± 0.4 | 0.63 ± 0.01 |
| **6** | 345.13 ± 0.01 | 69.1 ± 0.4 | 88.8 ± 0.5 | 0.78 ± 0.01 |

**Table S2. Regions of holo-AGT variants with changes in stability (Δ%$D_{av}$ ≥ |10|) upon NB-AGT binding.** Destab.- region destabilized; Stab.- region stabilized.

| AGT | NB-AGT |
|---|---|
| | **1** |
| **WT** | None |
| **LM** | Destab.: 27-43, 54-57, 204-209, 261-269. |
| **LM-G170R** | Destab.: 26-44, 54-57, 204-208, 244-248, |
| **Consensus** | **Destab.: 26-44, 54-57, 204-209, 244-248, 261-269.** |
| | **2** |
| **WT** | Stab.: 173-178. |
| **LM** | Stab.: 173-178. |
| **LM-G170R** | Stab.: 173-178. |
| **Consensus** | **Stab.: 173-178.** |
| | **3** |
| **WT** | Stab.: 105-110, 123-136, 231-234, 349-357. |
| **LM** | Stab.: 105-110, 123-134, 231-234, 348-357. |
| **LM-G170R** | Stab.: 105-111, 123-128, 231-234, 347-357. |
| **Consensus** | **Stab.: 105-111, 123-136, 231-234, 347-357.** |
| | **4** |
| **WT** | Stab.: 102-111, 121-134, 231-234, 251-254, 347-359. |
| **LM** | Stab.: 105-111, 121-128, 231-234, 250-255, 347-359. |
| **LM-G170R** | Stab.: 104-111, 121-128, 231-234, 250-255, 347-359. |
| **Consensus** | **Stab.: 102-111, 121-134, 231-234, 347-359.** |
| | **5** |
| **WT** | Stab.: 102-112, 123-136, 231-234. |
| **LM** | Stab.: 104-111, 123-128, 231-234. |
| **LM-G170R** | Stab.: 102-111, 123-128, 231-234. |
| **Consensus** | **Stab.: 102-112, 123-136, 231-234.** |
| | **6** |

| | |
|---|---|
| **WT** | Stab.: 302, 303-309, 310, 326-334, 381-392. |
| **LM** | Stab.: 302, 303-309 , 310, 327-330, 384-392. |
| **LM-G170R** | Stab.: 302, 303-309, 310, 324-330, 383-392 |
| **Consensus** | **Stab.: 302-310, 324-330, 382-392.** |

**Table S3. Kinetic parameters of AGT-Ma and G170R-Mi in the absence and presence of NB-AGT-3 at two different concentrations.** The kinetic parameters were measured in potassium phosphate 0.1 M pH 7.4 for all the conditions.

| AGT variant | [NB-AGT-3](μM) | $k_{cat}$ ($s^{-1}$) | $K_M$ (L-Ala) (mM) | $k_{cat}/K_M$ ($s^{-1}/mM^{-1}$) |
|---|---|---|---|---|
| WT | 0 | 44 ± 2 | 24 ± 2 | 1.8 ± 0.2 |
| | 1 | 45 ± 3 | 26 ± 3 | 1.7 ± 0.2 |
| | 100 | 42 ± 2 | 24 ± 3 | 1.7 ± 0.2 |
| LM-G170R | 0 | 33 ± 4 | 22 ± 4 | 1.4 ± 0.3 |
| | 1 | 34 ± 2 | 28 ± 5 | 1.2 ± 0.2 |
| | 100 | 32 ± 4 | 30 ± 7 | 1.0 ± 0.3 |
